# Evolved resistance to GAPDH inhibition results in loss of the Warburg Effect but retains a different state of glycolysis

**DOI:** 10.1101/602557

**Authors:** Maria V. Liberti, Annamarie E. Allen, Vijyendra Ramesh, Ziwei Dai, Katherine R. Singleton, Zufeng Guo, Jun O. Liu, Kris C. Wood, Jason W. Locasale

## Abstract

Aerobic glycolysis or the Warburg Effect (WE) is characterized by increased glucose uptake and incomplete oxidation to lactate. Although ubiquitous, the biological role of the WE remains controversial and whether glucose metabolism is functionally different during fully oxidative glycolysis or during the WE is unknown. To investigate this question, we evolved resistance to koningic acid (KA), a natural product shown to be a specific inhibitor of glyceraldehyde-3-phosphate dehydrogenase (GAPDH), a rate-controlling glycolytic enzyme during the WE. We find that KA-resistant cells lose the WE but conduct glycolysis and surprisingly remain dependent on glucose and central carbon metabolism. Consequentially this altered state of glycolysis leads to differential metabolic activity and requirements including emergent activities in and dependencies on fatty acid metabolism. Together, these findings reveal that, contrary to some recent reports, aerobic glycolysis is a functionally distinct entity from conventional glucose metabolism and leads to distinct metabolic requirements and biological functions.

## INTRODUCTION

Glycolysis, the uptake and metabolism of glucose, is a set of chemical reactions conserved to the most primitive of organisms and is fundamental to sustaining life (Bar-Even et al., 2012). Together with the catabolism of amino acids and lipids, glycolysis is a component of central carbon metabolism. The breakdown of glucose to pyruvate allows for the generation of energy in the form of ATP and reducing equivalents. In addition, glucose can be diverted into biosynthetic pathways for anabolic metabolism and for the process of metabolizing glucose to confer signaling functionality by, for example, coupling to the generation of reactive oxygen species and the mediation of chromatin state (Chandel, 2015).

Given the numerous biological functions that are conferred through glycolysis, it is reasonable to speculate that glucose metabolism could exist in a number of distinct phenotypically defined states. For example, aerobic glycolysis or the Warburg Effect (WE) (increased glucose uptake and incomplete oxidation to lactate in the presence of oxygen) is canonically thought to constitute a switch from fully oxidative glycolysis, the more common form of metabolism observed in differentiated cells (DeRisi et al., 1997; Vander Heiden et al., 2009). This altered form of glucose metabolism has been thought to result in differential functionality in cells including variations in anabolic metabolism.

While aerobic glycolysis has been extensively studied, there are contradictions regarding the biological function that it confers in cells separate from glucose metabolism. For example, findings from recent studies have questioned whether a switch from mitochondrial metabolism does in fact occur during the WE (Sellers et al., 2015) which is consistent with original observations (Crabtree, 1929; Warburg et al., 1927). Furthermore, other work has challenged whether the WE has any specific role in anabolic metabolism (DeBerardinis et al., 2007; Hosios et al., 2016). Indeed, studies have defined metabolic requirements of cancer to exist in two states – glycolysis-dependent or independent (Boudreau et al., 2016; Pusapati et al., 2016). If this is the case, then the WE would not be biologically distinct from other aspects of glucose metabolism. Thus, there is a surprising lack of clarity in the current literature as to whether the WE is a biologically defined state (Liberti and Locasale, 2016).

Metabolic control analysis has indicated that GAPDH exerts more control over glycolysis during the WE (Bakker et al., 2000; Shestov et al., 2014) and thus partial inhibition of GAPDH has a large differential effect on reducing glycolysis in WE conditions (Liberti et al., 2017). As a proof of concept, a natural product, koningic acid (KA), was shown to be selective for GAPDH as expression of a resistant allele of GAPDH ablates all changes in metabolism induced by the compound (Kumagai et al., 2008; Liberti et al., 2017; Watanabe et al., 1993). The compound is also selective against the fitness of cells specifically undergoing aerobic glycolysis (Kornberg et al., 2018; Liberti et al., 2017). Furthermore, numerous studies have suggested that targeting of GAPDH may be beneficial (Ganapathy-Kanniappan et al., 2009; Louie et al., 2016; Yun et al., 2015), in retrospect through partial inhibition of GAPDH during aerobic glycolysis. Thus, KA is a valuable probe to study the WE and to determine whether it has any specific biological function outside of glycolysis.

In this study, we sought to address the question of whether the WE can be phenotypically defined apart from glycolysis and fully oxidative glucose metabolism. Using a series of pharmacological and metabolomic approaches, we provide evidence that glucose metabolism exists in a number of defined metabolic states. Using acquired resistance to GAPDH inhibition as a model and KA as a tool, we show that cells can simultaneously evolve loss of the WE but continue to remain dependent on glycolysis. Consequently, these cells that have a selection pressure to lose the WE display widespread changes in metabolism downstream of glycolysis changes including a marked rewiring of fatty acid metabolism. Thus, our study provides evidence that the WE can be a biologically distinct form of glycolysis.

## RESULTS

### GAPDH inhibition leads to different outcomes from targeting glucose uptake

We first sought to determine whether disrupting GAPDH activity results in different outcomes from other perturbations to glycolysis. Since GAPDH has differential rate control in cells undergoing the WE (i.e. high glucose uptake and lactate secretion) (Liberti et al., 2017), we used a high WE cell line, BT-549, and compared inhibition of GAPDH with KA to inhibition of glucose uptake and deprivation of glucose from the culture media (Figure 1A). First, we measured the IC50 of E11, a validated, highly potent inhibitor of GLUT-1 and thus glucose uptake (Liu et al., 2017) (Figure S1A), and compared cell viability of BT-549 treated with doses of KA above and below the known IC50 (Liberti et al., 2017) and/or E11. We found that co-treatment of KA and E11 caused more significant decreases in cell viability than with either compound alone (Figure 1B). Given these data, we carried out a dose-response assay with calculated combination indices (CIs) for E11 and KA in BT-549 as well as in other cell lines that have been previously reported to undergo the WE (Liberti et al., 2017) (Figures S1B-S1F). These results revealed synergy between KA and E11 at low/middle concentrations of KA, and also highlighted differential effects of GAPDH and GLUT-1inhibition on cell viability.

**Figure 1.**
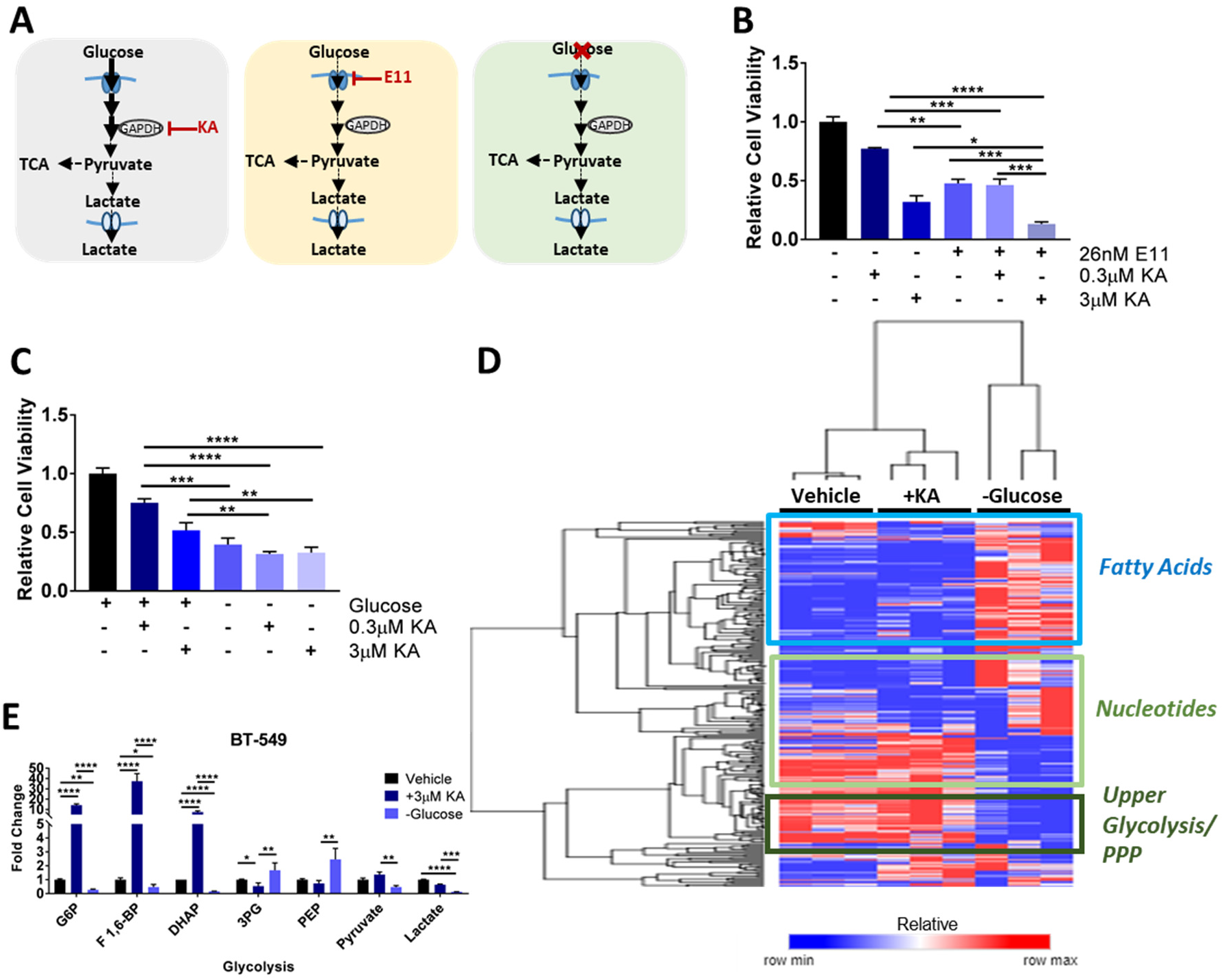
GAPDH inhibition leads to different outcomes from targeting glucose uptake. **(A)** Schematic representing the comparison of KA treatment to glucose transporter-1 (GLUT-1) inhibition with E11 or deprivation of glucose from the growth media. **(B)** Cell viability of BT-549 breast cancer cells treated with E11 (26nM) with or without KA (0.3μM or 3μM) after 24 hours. *p<0.05, **p<0.01, ***p<0.001, ****p<0.0001 as determined by One-Way ANOVA. **(C)** Cell viability of BT-549 breast cancer cells cultured in complete or glucose restricted media and treated with or without KA (0.3μM or 3μM) after 24 hours. *p<0.05, **p<0.01, ***p<0.001, ****p<0.0001 as determined by One-Way ANOVA. **(D)** Hierarchical clustered heatmap quantile normalized of BT-549 cells with condition annotations of global metabolic responses to vehicle, 3μM KA, or glucose restricted conditions for 6 hours with annotations of metabolic pathways. Scale represents 0 to 1 for row min and row max respectively after quantile normalization. **(E)** BT-549 glycolysis profile for vehicle, KA, or glucose restricted conditions for 6 hours. G6P, glucose-6-phosphate; F 1,6-BP, fructose 1,6-bisphosphate; DHAP, dihydroxyacetone phosphate; 3PG, 3-phosphoglycerate; PEP, phosphoenolpyruvate. All data are represented as mean ± SEM from n=3 biological replicates. *p<0.05, **p<0.01, ***p<0.001, ****p<0.0001 as determined by Two-Way ANOVA unless otherwise indicated.

In line with these findings, we further found significant changes between KA treated cells and cells cultured in glucose-deprived media (Figures 1C), as well as synergy between KA and glucose deprivation at low/middle concentrations of KA (Figures S1G-S1J). To assess these differential effects on cells at a metabolic level, we used liquid chromatography coupled to high resolution mass spectrometry (LC-HRMS) based metabolomics which revealed gross differences in global metabolism when comparing KA treated BT-549 cells to glucose-deprived cells (Figure 1D). An analysis of glycolysis indicated an accumulation of glycolytic intermediates upstream and depletion of those downstream of GAPDH in cells treated with KA, whereas glucose-deprived conditions revealed an overall depletion of metabolites throughout glycolysis (Figure 1E). Thus, cells exhibit a differential response to KA compared to other modes of glycolysis inhibition. Together, these data provide rationale that glycolysis could exist in multiple states given the differences seen in global metabolic and phenotypic responses upon disruption of the pathway at different steps, which we sought to test further.

### Cells evolve resistance to GAPDH inhibition independent of drug metabolism

To investigate the possibility that glycolysis could exist in different biological states, we hypothesized that cells could transition from the WE to another state of glucose metabolism when faced with a selective pressure against maintaining glycolysis in a certain state. Given that KA was previously shown to be selectively toxic to cells undergoing the WE, we suspected that it could be a useful tool to investigate this concept. We cultured BT-549 cells with incrementally increasing concentrations of KA and monitored their growth rate over a period of 20 weeks with parental cells maintained in cell culture in parallel (Figures 2A and 2B). Once cells developed resistance to 3μM KA, three clonal cell populations were isolated and maintained in 3μM KA for the remainder of the study (BT-549(R)1-3) (Figure 2C). IC50 values for KA in each of these clones were found to be greater than 200μM compared to the parental cells that exhibited an IC50 of ~1μM KA (Figure 2D).

**Figure 2.**
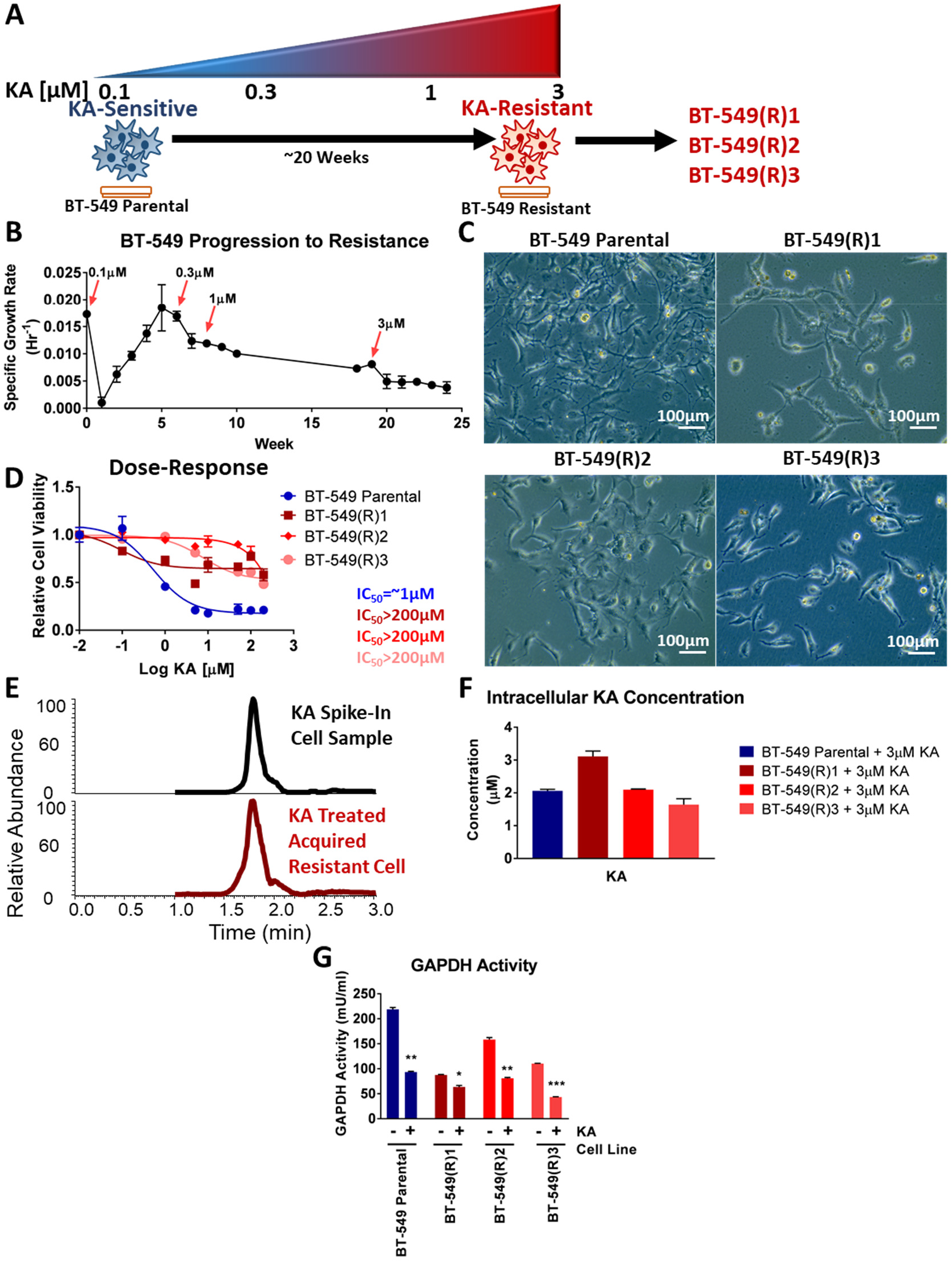
Cells evolve resistance to GAPDH inhibition independent of drug metabolism. **(A)** Schematic representing progression to acquired resistance model of BT-549 cells. After 20 weeks of incrementally increasing doses of KA clonal populations of acquired resistant cells were isolated and maintained in 3μM KA for the duration of the study. **(B)**Recorded growth rates of BT-549 cells during the 20-week period of progression to acquired resistance. **(C)** Representative images of BT-549 parental (top left) and 3 clonal acquired resistant cells (top right, bottom row). **(D)** Cell viability of BT-549 parental and acquired resistant cells treated with 0-200μM KA and reported IC50 values. **(E)** Mass spectra of KA in a spike-in sample and KA treated acquired resistant cell using liquid chromatography-mass spectrometry (LC-MS) with a representative sample. **(F)** Intracellular KA concentrations from BT-549 parental and acquired resistant cells treated with 3 μM KA. **(G)** Relative GAPDH activity in BT-549 parental and acquired resistant cells in response to vehicle or 3μM KA (n=2). All data are represented as mean ± SEM from n=3 biological replicates unless otherwise indicated. *p<0.05, **p<0.01, ***p<0.001, ****p<0.0001 as determined by multiple t-tests.

For this system to be an effective model for evolving a transition out of the WE, it was necessary to first to determine whether resistance to KA was occurring due to mechanisms outside of cellular metabolism. There are several known pharmacological mechanisms that are commonly implicated in drug resistance that include alterations in drug metabolism and target disengagement (Holohan et al., 2013). To determine whether resistance to KA developed due to altered drug metabolism such as a difference in drug efflux, we used LC-HRMS to measure intracellular concentrations of KA in BT-549 sensitive and acquired resistant cells as well as in MCF-7 KA-intrinsic resistant cells (Figure 2E). Intracellular concentrations of KA in BT-549 acquired resistant cells remained consistent with concentrations detected in BT-549 parental and MCF-7 intrinsic resistant cells that were treated with 3μM KA (Figures 2F and S2A). We also found similar intracellular concentrations of KA between BT-549 parental and acquired resistant cells at different timepoints (Figures S2B-S2D).

To determine whether KA was still engaging its target, the catalytic site of GAPDH (Endo et al., 1985; Sakai et al., 1988), we carried out a GAPDH activity assay in the presence or absence of KA and found that KA maintains target engagement through decreasing GAPDH activity in BT-549 acquired resistant cells comparable to BT-549 parental cells (Figures 2G, S2E and S2F) and MCF-7 cells (Figure S2G). In support of these findings, we observed little difference in GAPDH protein expression between BT-549 parental and acquired resistant cells (Figure S2H). We also found that acquired resistant cells can become re-sensitized to KA upon KA removal for two weeks, followed by addition of KA again (Figure S2I), further arguing against an inability to maintain target engagement. Together, these data suggest that acquired resistant cells retain normal KA drug metabolism properties with continued target engagement at the active site of GAPDH. Thus, independent of drug pharmacology, biological mechanisms related to glucose metabolism may underlie the resistance to GAPDH inhibition.

### Acquired resistant cells remain dependent on glycolysis, but lose the Warburg effect

Given that these cells evolved resistance to the glycolytic enzyme GAPDH, we sought to determine whether they continued to depend on glycolysis for survival. We treated BT-549 acquired resistant cells with KA and found that acquired resistant cells still undergo glycolysis marked by increases in fructose 1,6-bisphosphate (F 1,6-BP) compared to untreated BT-549 parental, which are indicative of slower rates of glycolysis (Shestov et al., 2014; van Heerden et al., 2014) and decreases in lactate levels, suggestive of a lower WE (Figure 3A). Interestingly, we also found that acquired resistant cells exhibited similar or lower levels of pentose phosphate pathway metabolites compared to parental cells (Figure S3A-S3D).

**Figure 3.**
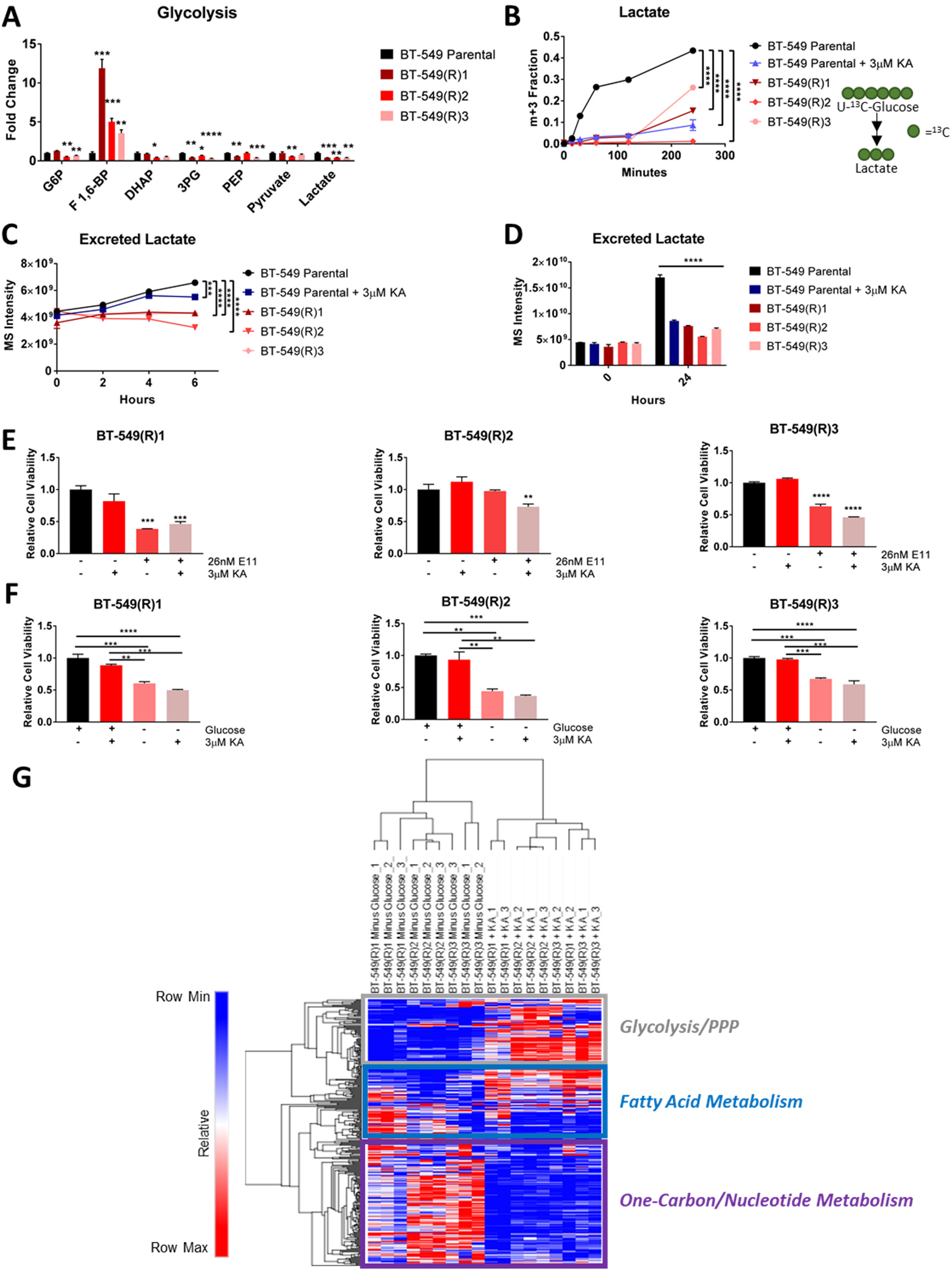
Acquired resistant cells remain dependent on glycolysis, but lose the Warburg Effect. **(A)** Glycolytic metabolite levels. *p<0.05, **p<0.01, ***p<0.001, ****p<0.0001 as determined by multiple t-tests. **(B)** ^13^C-lactate from (U-^13^C)-glucose in BT-549 parental treated with vehicle or 3μM KA and acquired resistant cells for 0-4 hours. **(C)** Excreted lactate detected in media from 0-6 hours. **(D)** Excreted lactate detected in media after 24 hours. **(E)** Cell viability of BT-549 acquired resistant cancer cells treated with E11 (26nM) with or without KA (3μM) after 24 hours. *p<0.05, **p<0.01, ***p<0.001, ****p<0.0001 as determined by One-Way ANOVA. **(F)** Cell viability of BT-549 acquired resistant cancer cells cultured in complete or glucose restricted media and treated with or without KA (3μM) after 24 hours. *p<0.05, **p<0.01, ***p<0.001, ****p<0.0001 as determined by One-Way ANOVA. **(G)** Hierarchical clustered heatmap quantile normalized of BT-549 cells with condition annotations of global metabolic responses to vehicle, 3μM KA, or glucose restricted conditions for 6 hours with annotations of metabolic pathways. Scale represents 0 to 1 for row min and row max respectively after quantile normalization. G6P, glucose-6-phosphate; F 1,6-BP, fructose 1,6-bisphosphate; DHAP, dihydroxyacetone phosphate; 3PG, 3-phosphoglycerate; PEP, phosphoenolpyruvate. All data are represented as mean ± SEM from n=3 biological replicates. *p<0.05, **p<0.01, ***p<0.001, ****p<0.0001 as determined by Two-Way ANOVA unless otherwise indicated.

Given the low lactate levels in the acquired resistant cells, we next sought to examine lactate production from glucose using uniformly labeled (U-^13^C)-glucose in BT-549 parental and acquired resistant cells. We found decreased lactate production in the acquired resistant cells compared to untreated parental cells at levels comparable to the lactate production observed in the parental cells treated with 3μM KA (Figures 3B and S3E). In addition, we measured intracellular glucose and lactate levels in parental and acquired resistant cells (Figure S3F). As expected, parental and acquired resistant cells exhibited similar intracellular glucose levels, indicative of similar glucose uptake. Parental and acquired resistant cells had similar intracellular lactate levels, which we reasoned to likely be due to the majority of lactate produced by cells being immediately secreted out of cells and into the environment (Liberti and Locasale, 2016; Vander Heiden et al., 2009). Indeed, upon measuring excreted lactate from cells in spent media from 0-6 hours, we found significantly less excreted lactate from BT-549 acquired resistant cells compared to parental cells treated with either vehicle or KA, with a similar finding after 24 hours (Figures 3C, 3D, and S3G). In addition, the decreases in lactate levels were not immediately reversible upon removal of KA (Figure S3H). Quantitative measurements of the WE via relative lactate flux calculations revealed lower lactate flux (i.e. low WE) in acquired resistant cells compared to parental cells, with a lactate flux value comparable to the low glycolytic cell line, MCF-7 (Figure S3I). Together, these data indicate that acquired resistant cells evolved a loss of the WE.

Since acquired resistant cells no longer undergo the WE, we asked whether they remained dependent on glucose uptake, since previous studies have indicated that cells are either glycolysis-dependent or independent (Boudreau et al., 2016; Pusapati et al., 2016). We treated the acquired resistant cells with E11 and/or KA and determined cell viability. While these cells remained resistant to KA as expected, we surprisingly observed differences in sensitivity upon treatment with E11 in combination with KA (Figure 3E). To further investigate this differential dependence on glycolysis, we studied the response to glucose deprivation. After 24 hours of culture in glucose-deprived growth media and KA, we found that the viability of BT-549 acquired resistant cells decreased compared to cells cultured in full growth media and maintained in KA (Figure 3F). In addition, metabolite profiling of BT-549 acquired resistant cells in glucose-deprived growth media compared to those maintained in 3μM KA revealed global differences in overall metabolic levels (Figure 3G). Together, these data indicate that BT-549 acquired resistant cells remain dependent on glycolysis for survival but no longer exhibit or require the WE.

We further found that acquired resistant cells displayed decreased fraction labeling and carbon contribution from glucose through glycolysis, decreased fraction labeling with a small increase in carbon contribution from glucose through the pentose phosphate pathway, and decreased fraction labeling and carbon contribution through the citric acid (TCA) cycle (Figures S3J-S3T). Taken together, these data further clarify the existence of multiple states of glucose metabolism.

### Changes in fatty acid metabolism emerge as a functional output of evolved resistance to KA

Our data thus far indicate a transition from the utilization of the WE to loss of the WE in acquired resistant cells to KA. To understand the temporal dynamics of metabolism that result with the observed change in state of glucose metabolism, we extracted metabolites from cells at different times over the 20-week time course during progression to resistance (Figure 4A). Metabolite profiling revealed global differences with glycolysis most prominently affected in the parental cells, and changes to fatty acid, one-carbon and nucleotide metabolism most apparent in acquired resistant cells (Figures 4B and 4C). Over half of the common changes in each of the clones relative to the parental cells were related to fatty acid metabolism (Figure 4D), including small and variably significant increases in fatty acid levels (Figure S4A-S4C).

**Figure 4.**
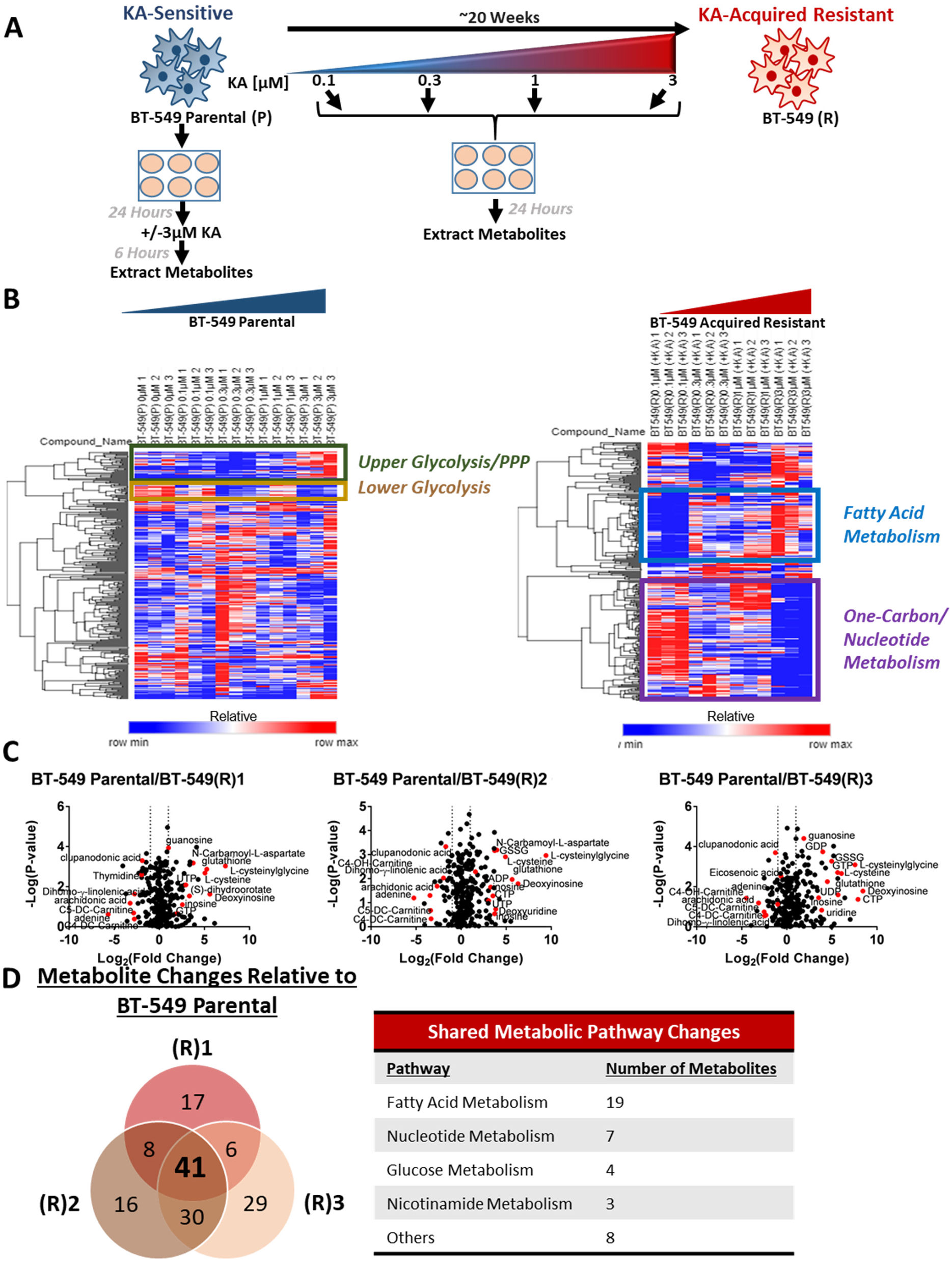
Changes in fatty acid metabolism emerge as a functional output of evolved resistance to KA. **(A)** Schematic of experimental setup for metabolomics during evolution to acquired resistance to KA. **(B)** Hierarchical clustered heatmap quantile normalized of 0-3μM KA dose-response in BT-549 parental and 0-3μM evolved resistance in BT-549 acquired resistant cells with annotations of metabolic pathways. Scale represents 0 to 1 for row min and row max respectively after quantile normalization. **(C)** Volcano plots showing metabolite profiles of BT-549 acquired resistant cells maintained in 3μM KA compared to BT-549 parental cells treated with vehicle. Log_2_ fold change versus –log_10_ p-value. Dotted lines along x-axis represent ± log_2_(1) fold change and dotted line along y-axis represents –log_10_(0.05). Metabolites ± log_2_(1) fold change shown as red points with metabolite names denoted. All other metabolites are black points. **(D)** Venn diagram indicating the overlap of metabolic changes among KA-resistant clones based on average ±log_2_(1) fold changes compared to BT-549 parental cells treated with vehicle. G6P, glucose-6-phosphate; F 1,6-BP, fructose 1,6-bisphosphate; DHAP, dihydroxyacetone phosphate; 3PG, 3-phosphoglycerate; PEP, phosphoenolpyruvate. All data are represented as mean ± SEM from n=3 biological replicates. *p<0.05, **p<0.01, ***p<0.001, ****p<0.0001 as determined by Two-Way ANOVA.

Given the common changes to fatty acid metabolism and our findings that acquired resistant cells are utilizing glucose carbon differently compared to parental cells, we asked whether other carbon sources, such as glutamine, account for the observed differences (Figure S4D). Interestingly, we found small but significant increases in the fraction labeled from glutamine through the TCA cycle indicating that acquired resistant cells fuel their TCA cycle, at least in part, by glutamine (Figures S4E-S4I). Next, we analyzed whether partial increases in fatty acid levels in acquired resistant cells result from carbon contribution by glucose, glutamine, and/or palmitate and found small contributions of each of these carbon sources into representative fatty acid metabolites (Figures S4J-S4Q). We next asked whether removal of lipids from the extracellular environment differentially affects acquired resistant cell survival (Figure S4R). We found no significant differences compared to parental cells, likely due to extracellular lipids not being a major carbon contributor in acquired resistant cells (Figure S4S-S4U). Together, these data demonstrate differences in fatty acid metabolism in acquired resistant cells with variable contributions from different carbon sources compared to parental cells.

To further probe the differences we observed in fatty acid metabolism, we used cerulenin, a fatty acid synthase inhibitor that inhibits fatty acid oxidation by increasing malonyl-CoA levels (Hu et al., 2003; Loftus et al., 2000; Thupari et al., 2001), and found increased sensitivity in acquired resistant cells to compared to parental cells and also found no significant change in MCF-7 cells (Figures S5A-S5C). Additionally, the acquired resistant cells treated with cerulenin exhibited a differential metabolic response (Figures 5A and S5D). Next, we asked whether cerulenin differentially affected acyl-carnitine levels, signatures of fatty acid metabolism (Gao et al., 2018; Koves et al., 2008), in BT-549 acquired resistant cells compared to BT-549 parental cells. While we found that BT-549 parental cells exhibited few changes in acyl-carnitine levels upon co-treatment with KA and cerulenin, we found that when acquired resistant cells are maintained in KA, acyl-carnitines are elevated but upon co-treatment with cerulenin many of them significantly decrease (Figures 5B and S5E).

**Figure 5.**
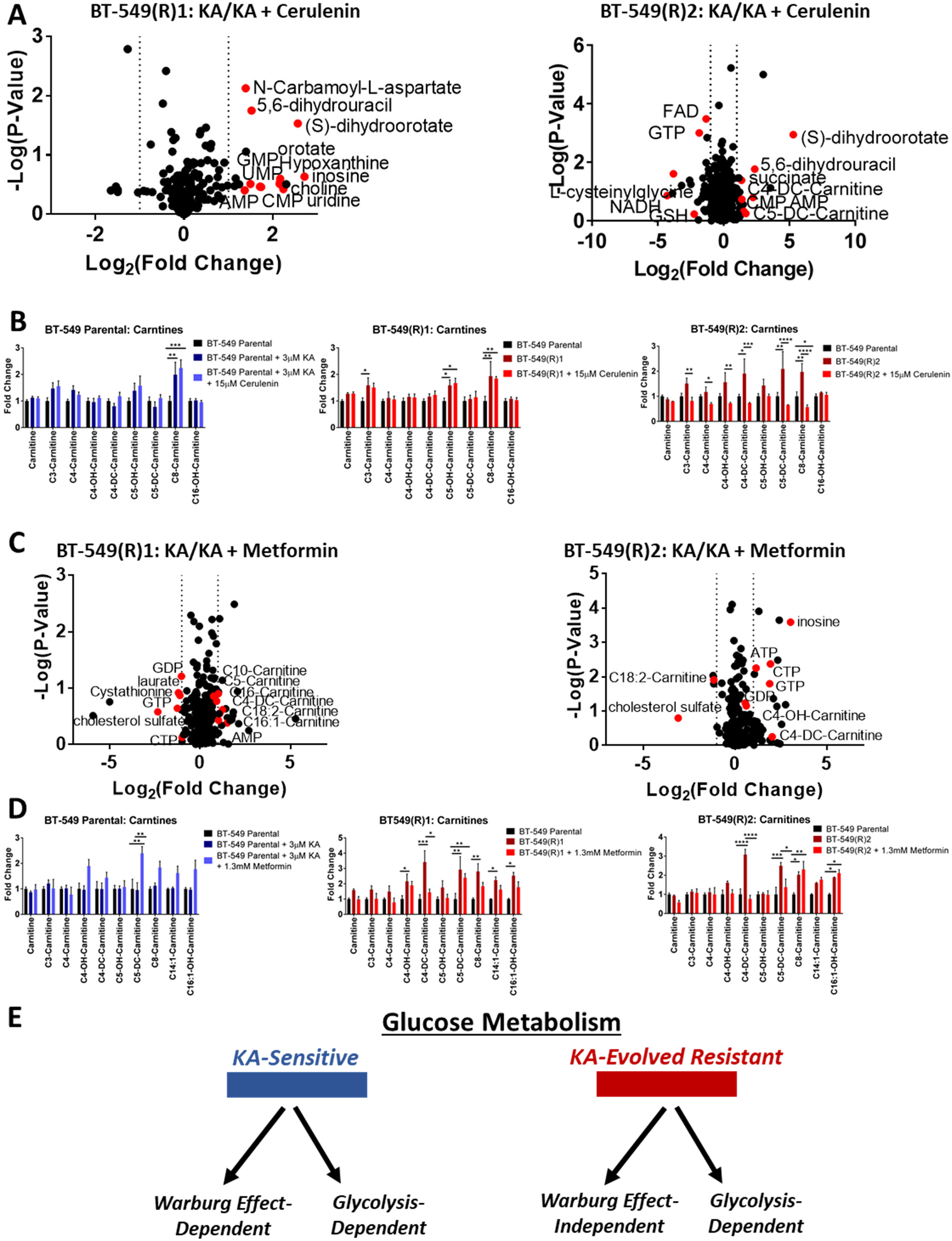
Changes in fatty acid metabolism occur downstream of differences in glycolysis in acquired resistance to KA. **(A)** Volcano plots showing metabolite profiles of BT-549 acquired resistant (R)1 and (R)2 cells maintained in KA (3μM) with or without cerulenin (15μM). Log_2_ fold change versus –log_10_ p-value. Dotted lines along x-axis represent ± log_2_(1) fold change and dotted line along y-axis represents –log_10_(0.05). Metabolites ± log_2_(1) fold change shown as red points with metabolite names denoted. All other metabolites are black points. **(B)** Acyl-carnitine levels in BT-549 parental and acquired resistant cells maintained in KA (3μM) and treated with or without cerulenin (15μM) for 6 hours. **(C)** Volcano plots showing metabolic profiles of BT-549 acquired resistant (R)1 and (R)2 cells maintained in KA (3μM) with or without metformin (1.3mM) as in (A). **(D)** Acyl-carnitine levels in BT-549 parental and acquired resistant cells maintained in KA (3μM) and treated with or without metformin (1.3mM) for 6 hours. **(E)** Schematic representing different phenotypically-defined glucose metabolism states. GMP, guanosine monophosphate; UMP, uridine monophosphate; AMP, adenosine monophosphate; CMP, cytidine monophosphate; FAD, flavin adenine dinucleotide; NADH, nicotinamide adenine dinucleotide; GSH, reduced glutathione; IMP, inosine monophosphate; CTP, cytidine triphosphate; SAH, s-adenosyl-l-homocysteine. All data are represented as mean ± SEM from n=3 biological replicates. *p<0.05, **p<0.01, ***p<0.001, ****p<0.0001 as determined by Two-Way ANOVA.

Since the biguanide metformin has been shown to decrease fatty acid and mitochondrial metabolism (Owen et al., 2000), we asked whether metformin displayed similar effects to cerulenin. Using the measured IC50 of metformin in BT-549 parental cells (Figure S5F), we cotreated parental and acquired resistant cells with metformin and KA, which revealed decreases in cell viability of both BT-549 sensitive cells and acquired resistant cells, albeit to a lesser extent in BT-549(R)2 cells and no significant response in the non-glycolytic MCF-7 cells (Figures S5G and S5H). We also found that upon co-treatment with KA and metformin compared to KA alone, acyl-carnitines were to a larger extent significantly decreased in acquired resistant cells than in parental cells (Figures 5C-5D and S5I-S5J), thus confirming that distinct metabolic phenotypes related to fatty acid metabolism occur downstream of the differences in glycolysis.

## DISCUSSION

### Evolved resistance to KA as a model to study the Warburg Effect

Previous work has shown that GAPDH has a specific regulatory role in aerobic glycolysis (Kornberg et al., 2018; Liberti et al., 2017; Shestov et al., 2014; Yun et al., 2015). Our current study extends from this understanding to show that cells which evolve resistance to a specific GAPDH inhibitor, KA, lose the WE but remain dependent on glycolysis with different metabolic outputs, thus demonstrating that glucose metabolism can exist in different states. In particular, we demonstrate that cells can exist in at least two separate states of glucose metabolism including WE-dependent and glycolysis-dependent or WE-independent and glycolysis-dependent (Figure 5E). Glycolysis has been thoroughly studied using models that ablate the expression of glycolytic enzymes or completely block the pathway (Christofk et al., 2008; Israelsen et al., 2013; Patra et al., 2013; Pusapati et al., 2016) which reduces overall or eliminates altogether the activity of glycolysis. Our model of evolved resistance to KA is useful to study glucose metabolism regulation without complete pathway inhibition. Instead, we were able to place a selection pressure against the fitness of using the WE during proliferation. Moreover, using metabolomics, we further demonstrate that the resulting pressure to lose the WE retains a requirement for glycolysis but alters several metabolic outputs in central carbon metabolism. For example, fatty acid metabolism is upregulated and selectively required in cells. Our extensive metabolomics analyses also provide insight on enzymatic activity changes. Based on our profiling, it is worth noting that many activities in enzymes have likely changed throughout glycolysis and branching pathways. Furthermore, we could capture the temporal dynamics of metabolic alterations from early to late timepoints of this evolution. In physiology, such as within a tumor microenvironment, it is likely that this range of metabolic plasticity allows for rapid adaptation to a dynamic environment.

### Glucose metabolism exists in distinct phenotypically-distinct states

We were thus able to study glycolysis under different configurations of metabolic activity as cells transition from the WE to another glycolytic state. We provide clear evidence for a distinction between the WE and glucose metabolism. Interestingly, previous literature and related drug development efforts have worked under the model that glycolysis functions as a binary switch (i.e. glucose dependent or independent) (Boudreau et al., 2016; Pusapati et al., 2016). In these cases, glucose dependence is identical to aerobic glycolysis. Our findings show that although KA-resistant cells no longer undergo the WE, they undergo glycolysis with less lactate production and remain dependent on glucose uptake. While the difference in intracellular lactate levels between parental cells treated with KA and acquired resistant cells maintained in KA were not as dramatic as was seen with measurements of secreted lactate, the majority of lactate produced by cells is immediately secreted into the environment (Liberti and Locasale, 2016; Vander Heiden et al., 2009), likely accounting for the larger differential found in excreted lactate. Thus, our data indicate that glucose metabolism exists functionally in a set of states. By evolving resistance to GAPDH inhibition with KA, we show that WE-undergoing cells that lose aerobic glycolysis do not simply switch to increased oxidative phosphorylation but maintain glucose metabolism in a separate biological state.

### The biology of and targeting the Warburg Effect

While aerobic glycolysis has been extensively studied over the years, whether the WE has a function aside from glycolysis has been challenged (Faubert et al., 2017; Hosios et al., 2016). Our findings provide evidence for the WE existing as a biologically functional state of glucose metabolism. From a therapeutic perspective, elucidating the WE as a distinct phenomenon from glucose metabolism provides rationale for continued efforts to target the WE while keeping all other forms of glycolysis intact. Such efforts are underway including various studies particularly focusing on the targeting of GAPDH (Kornberg et al., 2018; Louie et al., 2016; Yun et al., 2015). Previous studies already indicate the feasibility and tolerability of targeting GAPDH therapeutically (Kornberg et al., 2018; Liberti et al., 2017). Although this study by itself does not resolve whether the WE is in fact driving cancer or whether it is a metabolic consequence of cancer progression, our study does confirm that the WE is a real biological phenomenon with different biological and metabolic properties.

## Author Contributions

Conceptualization, MVL and JWL; Metabolomics, MVL; Cell Viability Assays, MVL, AEA, VR; Nutrient Restriction Assays, MVL; Data Analysis, MVL, AEA, VR, ZD, KRS, KCW; Provision of Essential Reagents, ZG and JOL; All Other Experiments, MVL; Writing, MVL and JWL; Supervision, MVL and JWL.

## Acknowledgements

We thank the members of the Locasale laboratory for their comments and advice. Support from the National Institutes of Health (R01CA193256 (JWL), R00CA168997 (JWL), F99CA222986 (MVL)), the National Science Foundation (DGE-1144153 (MVL)), and the Sloan Foundation (MVL) are gratefully acknowledged.

## METHODS

### Cell Culture

BT-549 and MCF-7 cells were cultured in full media containing RPMI-1640 (Gibco), 10% heat-inactivated fetal bovine serum (FBS), 100U/ml penicillin and 100μg/ml streptomycin. BT-549 and MCF-7 cells were obtained from the American Tissue Culture Collection (ATCC). Koningic acid (KA)-resistant BT-549 cells were cultured and maintained in full media containing RPMI-1640, 10% heat-inactivated FBS, 100U/ml penicillin,100μg/ml streptomycin, and 3μM KA (isolated in-house) (Liberti et al., 2017). Cells were cultured in a 37°C, 5% CO2 atmosphere.

### Time to Progression to Resistance Assay

Cells were allowed to progress to resistance as previously described (Singleton et al., 2017). To allow cells to acquire resistance to KA, BT-549 breast cancer cells were first seeded in triplicate in 15cm plates at 3×10^6^ cells per plate in normal media. After 24 hr, the normal growth media was replaced with fresh media at the indicated KA treatment. After seven days, cells were lifted with 0.25% trypsin (Cellgro) and counted using Moxi Z mini automated cell counter. All cells up to 1×10^6^ cells were centrifuged at 1,500 rpm for 3 min and resuspended in 10ml of media and plated into a 15 cm plate with fresh treatment. For each measurement, once cell number reached 3×10^6^ cells two weeks in a row, the dose was increased as indicated. This procedure was repeated weekly for 20 weeks. Weekly growth rates (μ) were calculated from the number of cells plated the previous week (N_0_) and the number of cells counted on the current week (N) according to the formula

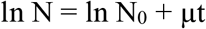

where t is elapsed time in hr. These growth rates were then used to project total cell number as if no cells had been discarded.

### Cell Viability Assays

For all cell lines, 5×10^4^ cells/well were seeded in triplicate in a 96-well plate and allowed to adhere for 24 hr. The following day, vehicle or treatment was added to each well at the respective concentrations. After 24 hr, the media was aspirated and replaced with 100μL phenol-red free RPMI-1640 and 12mM 3-[4,5-Dimethylthiazol-2-yl]-2,5-diphenyltetrazolium (MTT) (Thermo Fisher Scientific, #M6494) was added to the cells. After 4 hr, the media containing MTT was aspirated and 50 μL DMSO was added to dissolve the formazan and read at 540nm.

### Drug Treatments

For all cell lines, IC_50_ values of KA were measured by seeding 5×10^4^ cells/well in triplicate in a 96-well plate and allowed to adhere for 24 hr. The following day, media was changed and concentrations of either vehicle (H2O or DMSO), KA, E11 (Liu et al., 2017), cerulenin (Sigma-Aldrich, #C2389), or metformin (Santa Cruz Biotechnology, #202000A), were added. After 24 hr, cell viability assays were carried out using MTT as previously described.

### Nutrient Restriction in Media

For all cell lines, 5×10^4^ cells/well were seeded in triplicate with complete RPMI-1640 media in 96-well plates and allowed to adhere for 24 hr. On the following day, the respective treatment media was added in the absence or presence of KA at the indicated treatments. MTT assays were carried out as previously described. Treatment media used was as follows: Minus glucose (-Glucose): RPMI-1640 with glutamine lacking glucose containing 10% dialyzed FBS (Life Technologies), 100U/ml penicillin, and 100μg/ml streptomycin; Delipidated serum (D-FBS): RPMI-1640 with glutamine containing 10% delipidated FBS (ThermoFisher Scientific, #A3382101), 100U/ml penicillin, and 100μg/ml streptomycin. Cells were grown at 37°C with 5% CO_2_.

### Synergism Experiments

BT-549, SK-MEL-28, SK-MEL-5, and NCI-H522 cell lines were seeded at 5×10^4^ cells per well in 96-well plates and allowed to adhere for 24 hours prior to treatment. After, 0, 0.5, 1, 5, and 10μM KA was used as single concentrations or in combination with 0, 10, 100, 500, 1000nM E11 or in combination with 0, 0.5, 1, 5, 11mM glucose. After 24 hours, dose-response curves were generating using MTT reagent as described above. Then, cell viability values and concentrations with inputted into CompuSyn 1.0 software (https://compusyn.software.informer.com/) and combination indices (CIs) were calculated by the software to determine synergism, additivity, or antagonism.

### Re-sensitization of BT-549 acquired resistant cells to KA

KA was removed from BT-549 acquired resistant cells for 3 passages (~2 weeks) (BT-549(R)1-3^−KA,p.3^). After, parental cells, acquired resistant cells maintained in 3 μM KA, and BT-549(R)1-3^−KA,p.3^) cells were treated with vehicle, 0.5μM, or 3μM KA for 24 hours followed by measurement of cell viability with MTT reagent as previously described.

### GAPDH Activity Assay

GAPDH activity assay kit (BioVision, #K680) was used. All cells were seeded at 1×10^6^ cells per 10cm plate with either vehicle or KA. After 24 hr, cells were lysed, NADH standard curve was made, and cells were measured at 450nm in kinetic mode for 60 min at 37°C according to the manufacturer’s instructions.

### Microscopy

Cells were seeded at a density of 5×10^3^ cells per well in 6-well plates and allowed to adhere for 24 hours prior to treatment. After 48 hours, images were captured using a Leica DM IL LED microscope equipped with a Leica MC170HD camera at 10x objective using LAS EZ software (Leica). Scale bars = 100μm.

### Stable Isotope Labeling

Cells were seeded at 3×10^5^ cells/well in a 6-well plate and allowed to adhere for 24 hr. For U-^13^C-glucose isotopic labeling, cells were treated with either vehicle or KA for 6 hr, then replaced with RPMI-1640 media containing 11mM U-^13^C-glucose (Cambridge Isotope Laboratories, #CLM-1396) and vehicle or KA for 0-4 hr or for 24 hr. Metabolites were then extracted. For U-^13^C-glutamine isotopic labeling, cells were treated with either vehicle or KA for 6 hr, then replaced with RPMI-1640 media containing 10% dialyzed FBS, 2mM U-^13^C-glutamine (Cambridge Isotope Laboratories, #CLM-1822), and vehicle or KA for 24 hr. Metabolites were then extracted. For U-^13^C-palmitate isotopic labeling, cells were treated with vehicle or KA for 6 hr, then replaced with RPMI-1640 media containing 10% dialyzed FBS, 100μM U-^13^C-palmitiate (Cambridge Isotope Laboratories, #CLM-409) and vehicle or KA for 24 hr. Metabolites were then extracted.

### Intracellular Metabolite Measurements

Cells were seeded at 3×10^5^ cells/well in a 6-well plate and allowed to adhere for 24 hr. After, cells were treated with vehicle or KA for 6 hr and then washed twice with 0.9% NaCl. Metabolites were then extracted.

### Extracellular Metabolite Excretion Measurements

Cells were seeded at 3×10^5^ cells/well in a 6-well plate and allowed to adhere for 24 hr. After, cells were treated with vehicle or KA and 15μl media was collected from 0-4 hr and 24hr. Metabolites were then extracted.

### Metabolite Extraction

Metabolite extraction and subsequent Liquid Chromatography coupled to High Resolution Mass Spectrometry (LC-HRMS) for polar metabolites of each cell line was carried out using a Q Exactive Plus as previously described (Liu et al., 2014a; Liu et al., 2014b). For culture from adherent cell lines, media was quickly aspirated. Next, 1ml of extraction solvent (80% methanol/water) cooled to −80°C overnight was added immediately to each well and the plates were then transferred to −80°C for 15 min. After, the plates were removed, and cells were scraped into the extraction solvent on dry ice. For media extractions, 15μl media was collected at 0-24 hr. Next, 15μl extraction solvent (80% methanol/water) (optima LC-MS grade, Fisher Scientific, methanol, #A456; water, #W6) was added to the media. For absolute quantification of KA in cells, media was quickly aspirated, and cells were washed twice with 0.9% NaCl before following extraction for culture from adherent cells. 0.7μM KA in water was spiked into extraction solvent before centrifugation. For absolute quantification of KA at 2, 6, and 24 hr, cells were washed twice with 0.9% NaCl and a standard curve of KA was applied with concentrations from 0-25 μM KA spiked into methanol solvent of untreated BT-549 cells before centrifugation. All metabolite extractions were centrifuged at 20,000 *g* at 4°C for 10 min. Finally, the solvent in each sample was evaporated using a speed vacuum for metabolite analysis. For polar metabolite analysis, the cell metabolite extract was first dissolved in 15μl water, followed by dilution with 15 μl methanol/acetonitrile (1:1 *v/v*) (optima LC-MS grade, Fisher Scientific, methanol, #A456; acetonitrile, #A955). Samples were centrifuged at 20,000 *g* for 10 min at 4°C and the supernatants were transferred to LC vials. The injection volume for polar metabolite analysis was 5μl.

### Liquid Chromatography

An XBridge amide column (100 × 2.1 mm i.d., 3.5μm; Waters) was used on a Dionex (Ultimate 3000 UHPLC) for compound separation at room temperature. Mobile phase A is water with 5mM ammonium acetate, pH 6.9, and mobile phase B is 100% acetonitrile. The gradient is linear as follows: 0 min, 85% B; 1.5 min, 85% B; 5.5 min, 35% B; 10 min, 35% B; 10.5 min, 35% B; 10.6 min, 10% B; 12.5 min, 10% B; 13.5 min, 85% B; and 20 min, 85% B. The flow rate was 0.15 ml/min from 0 to 5.5 min, 0.17 ml/min from 6.9 to 10.5 min, 0.3 ml/min from 10.6 to 17.9 min, and 0.15 ml/min from 18 to 20 min. All solvents are LC-MS grade and purchased from Fisher Scientific.

### Mass Spectrometry

The Q Exactive Plus MS (Thermo Scientific) is equipped with a heated electrospray ionization probe (HESI) and the relevant parameters are as listed: evaporation temperature, 120°C; sheath gas, 30; auxiliary gas, 10; sweep gas, 3; spray voltage, 3.6 kV for positive mode and 2.5 kV for negative mode. Capillary temperature was set at 320°C, and S lens was 55. A full scan range from 70 to 900 (*m/z*) was used. The resolution was set at 70,000. The maximum injection time was 200 ms. Automated gain control (AGC) was targeted at 3×10^6^ ions.

### Peak Extraction and Data Analysis

Raw data collected from LC-Q Exactive Plus MS was processed on Sieve 2.0 (Thermo Scientific, https://www.thermofisher.com/order/catalog/product/IQLAAEGABSFAHSMAPV). Peak alignment and detection were performed according to the protocol described by Thermo Scientific. For a targeted metabolite analysis, the method “peak alignment and frame extraction” was applied. An input file of theoretical *m/z* and detected retention time of 197 known metabolites was used for targeted metabolite analysis with data collected in positive mode, while a separate input file of 262 metabolites was used for negative mode. *m/z* width was set to 10 ppm. The output file including detected *m/z* and relative intensity in different samples was obtained after data processing. If the lowest integrated mass spectrometer signal (MS intensity) was less than 1000 and the highest signal was less than 10,000, then this metabolite was considered below the detection limit and excluded for further data analysis. If the lowest signal was less than 1000, but the highest signal was more than 10,000, then a value of 1000 was imputed for the lowest signals. Mass isotopomer distributions (MID) were calculated and samples were normalized by comparing the ratio of glucose-derived labeled metabolites to unlabeled metabolites within each sample. Quantitation and statistics were calculated using Microsoft Excel and GraphPad Prism 7.0.

### Analysis of Metabolomics Data

GENE-E and Morpheus software were used for hierarchal clustering and heatmap generation (The Broad Institute, https://software.broadinstitute.org/GENE-E/index.html). For hierarchal clustering, spearman correlation parameters were implemented for row and column parameters, with the exception of BT-549 parental and acquired resistant drug response data, in which hierarchal clustering for row parameters only was used. Quantile normalization was used to normalize the data, represented by color scales.

### Lactate Flux Calculations

The time-dependent lactate labeling pattern was modeled as with the following equation:

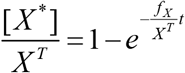

In which [*X*^*^] is the concentration of labeled lactate, *X^T^* is the total concentration (both labeled and unlabeled) of lactate, *f_X_* is the lactate production flux. This model was fit to lactate MIDs using the fit() function in MATLAB to determine relative lactate production fluxes. Relative lactate pool sizes were estimated from MS signal intensities.

### Quantification and Statistical Analysis

Unless otherwise noted, all error bars were reported ± SEM with n = 3 independent biological measurements and statistical tests resulting in p value computations were computed using a Student’s t test two tailed, multiple t-tests, one-way ANOVA. or two-way ANOVA of log transformed data followed by Tukey’s multiple comparisons. All statistics were computed using GraphPad Prism 7 (GraphPad, http://www.graphpad.com/scientific-software/prism/).

