## Supplemental Data for "Evolved resistance to GAPDH inhibition results in loss of the Warburg Effect but retains a different state of glycolysis"

### SUPPLEMENTARY FIGURES

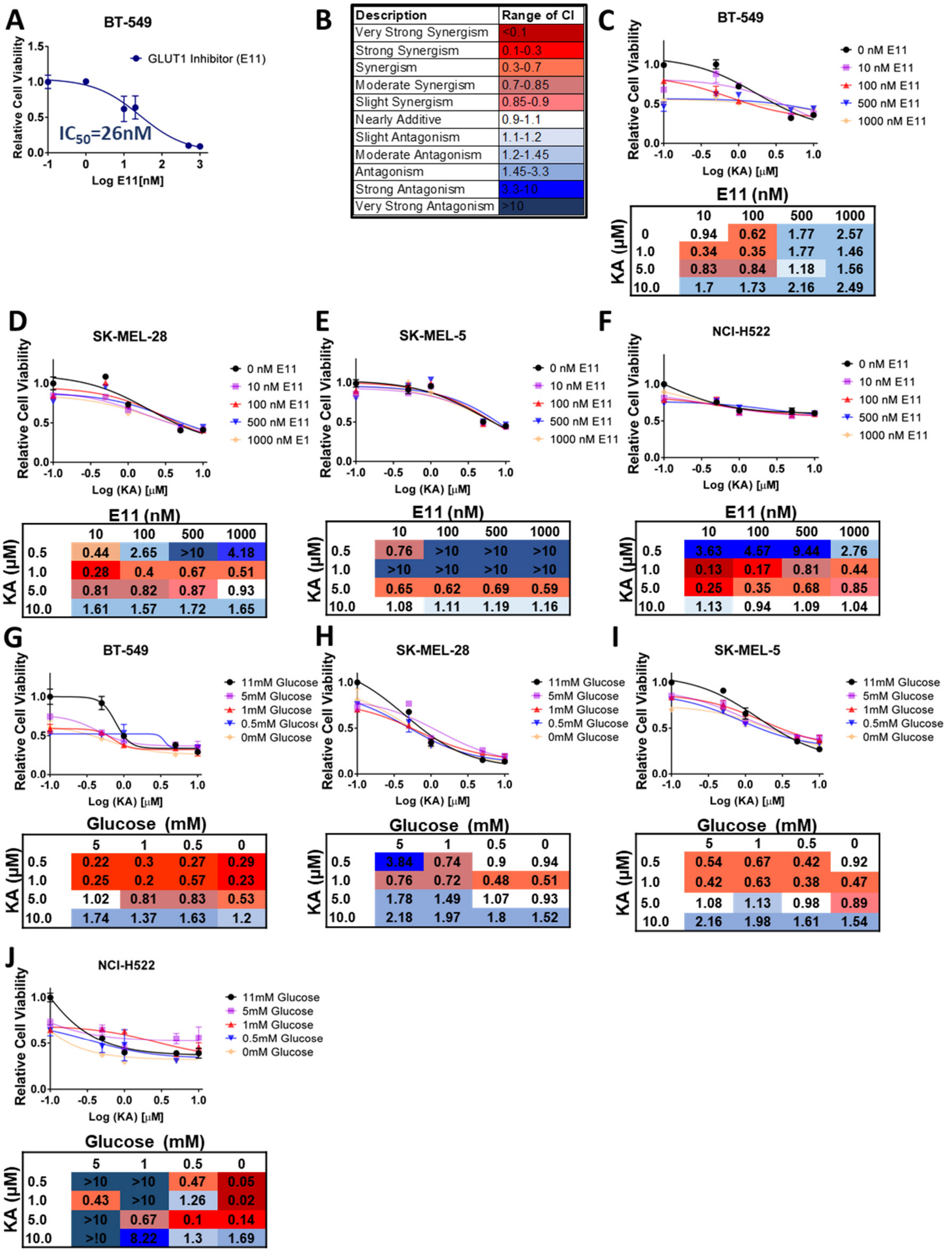


**Figure S1.** ***Related to Figure 1.*** **GAPDH inhibition leads to different outcomes from targeting glucose uptake.**

**(A)** BT-549 breast cancer cells dose-response curve after treatment with 0-1μM E11 for 24 hours.

**(B)** Description of synergy, additivity, and antagonism with respective combination indices (CI).

**(C)** Dose-response curve using 0-10μM KA and 0-1000nM E11 with associated calculated CIs in BT-549.

**(D)** SK-MEL-28 as in (C).

**(E)** SK-MEL-5 as in (C).

**(F)** NCI-H522 as in (C).

**(G)** Dose-response curve using 0-10μM KA and titration curve using 0-11mM glucose with associated calculated CIs in BT-549.

**(H)** SK-MEL-28 as in (G).

**(I)** SK-MEL-5 as in (G).

**(J)** NCI-H522 as in (G).

All data are represented as mean ± SEM from n=3 biological replicates.

*$p<0.05$, **$p<0.01$, ***$p<0.001$, ****$p<0.0001$ as determined Two-Way ANOVA.

CIs for synergy, additivity, and antagonism calculated using CompuSyn 1.0 software.

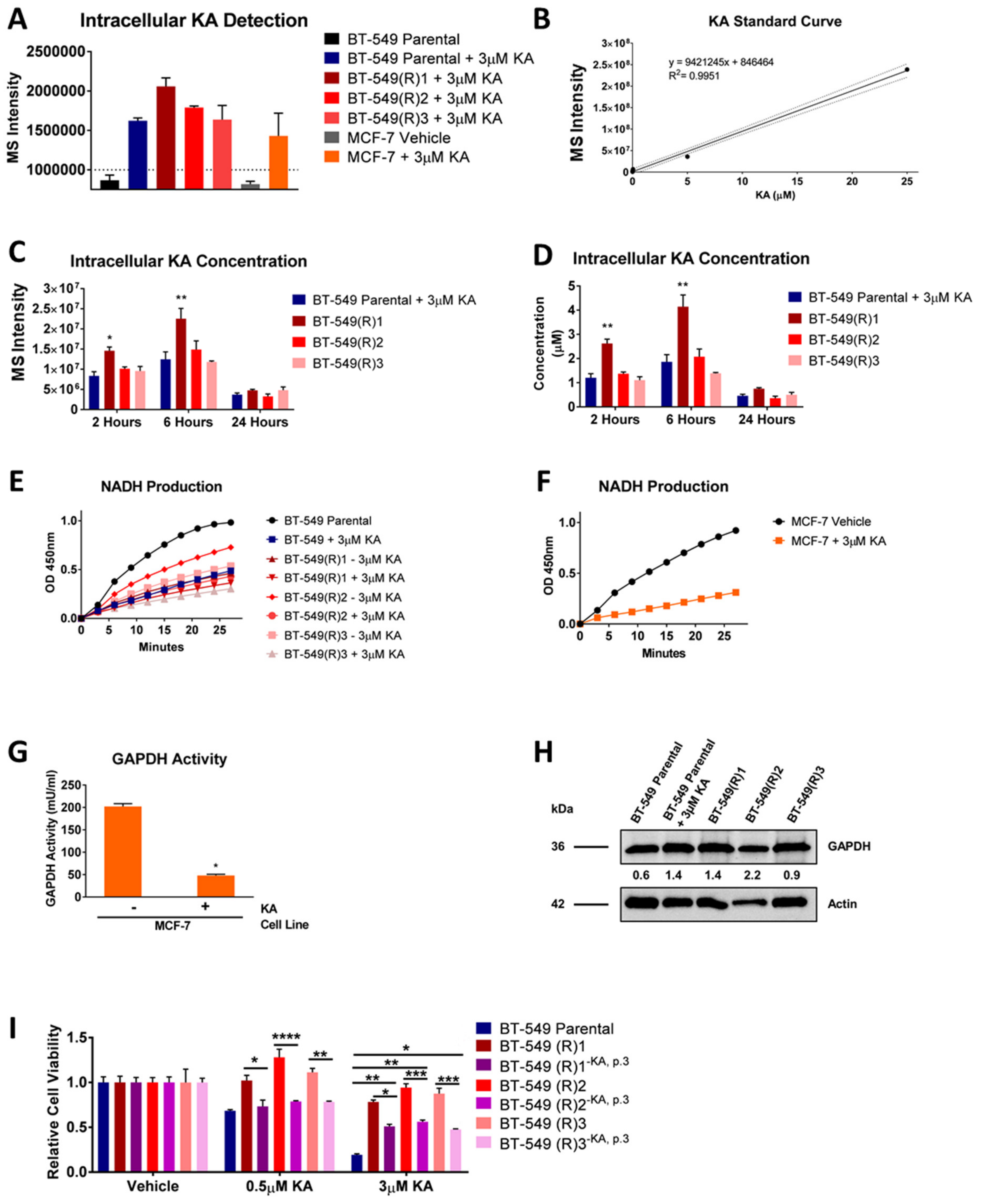


**Figure S2.** ***Related to Figure 2*. Cells evolve resistance to GAPDH inhibition independent of drug metabolism.**

**(A)** Mass spectrometry intensities of intracellular KA in vehicle or KA (3μM) treated cells using liquid chromatography-mass spectrometry (LC-MS). Dotted line denotes noise level.

**(B)** Standard curve of KA spiked into untreated BT-549 parental cells.

**(C)** MS intensity values of KA concentrations after treatment with vehicle or 3μM KA for 2, 6, and 24 hours.

**(D)** Absolute intracellular KA concentrations after treatment with vehicle or 3μM KA for 2, 6, and 24 hours.

**(E)** Relative NADH production from 0-27 minutes in kinetic mode of BT-549 parental and acquired resistant cells with and without KA (3μM) treatment (n=2).

**(F)** Relative NADH production of MCF-7 cells as in (B) (n=2).

**(G)** Relative GAPDH activity in MCF-7 cells in response to vehicle or 3μM KA (n=2). *p<0.05, **p<0.01, ***p<0.001, ****p<0.0001 as determined by student's t-test.

**(H)** Immunoblotting of GAPDH in BT-549 parental and acquired resistant cells with quantitation normalized to actin.

**(I)** BT-549 acquired resistant cells were passed either with KA or without KA (as indicated by superscript -KA, p.3) for 3 passages (a total of 2 weeks). After 2 weeks, cells were treated with vehicle, 0.5μM KA, or 3μM KA and compared to BT-549 parental cells.

All data are represented as mean ± SEM from n=3 biological replicates unless otherwise indicated.

*p<0.05, **p<0.01, ***p<0.001, ****p<0.0001 as determined by Two-Way ANOVA unless otherwise indicated.

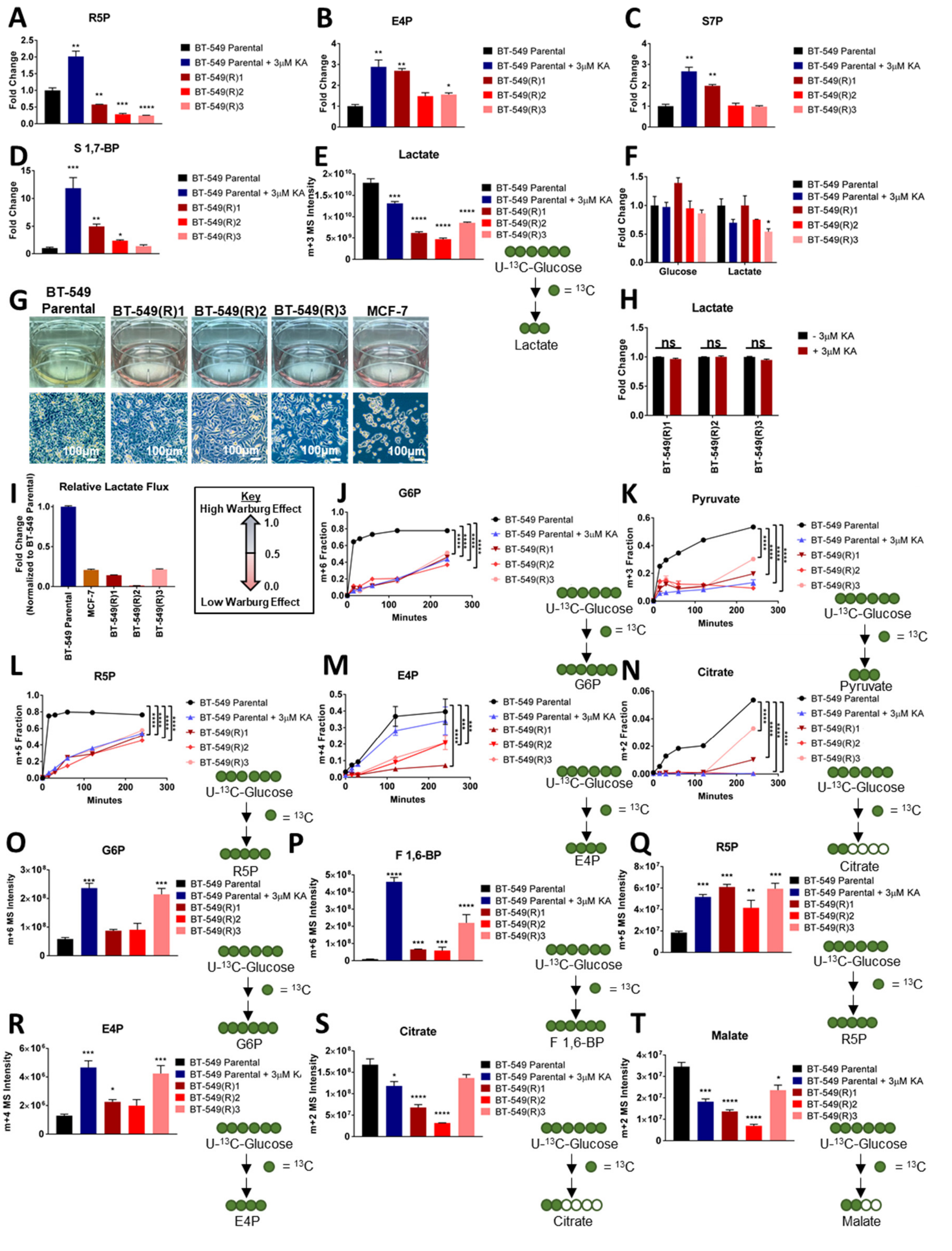

**Figure S3.** ***Related to Figure 3*. Acquired resistant cells remain dependent on glycolysis, but lose the Warburg Effect.**

**(A)** R5P levels in BT-549 parental and acquired resistant cells treated with vehicle or KA for 6 hours.

**(B)** E4P as in (A).

**(C)** S7P as in (A).

**(D)** S 1,7-BP as in (A).

**(E)** MS intensity of $^{13}$C-lactate in BT549 parental and acquired resistant cells treated with vehicle or KA for 6 hours followed by U-$^{13}$C-glucose labeling for 24 hours.

**(F)** Levels of intracellular glucose and lactate in BT-549 parental and acquired resistant cells after 6 hours.

**(G)** Representative images of spent media (top row) and cells (bottom row) at confluency in BT-549 parental, BT-549 acquired resistant, and MCF-7 intrinsic resistant cells after 48 hours.

**(H)** Lactate levels in BT-549 acquired resistant cells upon removal or maintained in KA (3μM) KA after 24 hours. Not significant denoted as "ns" determined by multiple t-tests.

**(I)** Relative lactate flux calculated for BT-549 parental and acquired resistant cells as well as in MCF-7 cells. Value are normalized to lactate flux of BT-549 parental cells. Key denotes low to high Warburg Effect and respective values.

**(J)** $^{13}$C-G6P in BT-549 parental and acquired resistant cells treated with vehicle or KA for 0-4 hours.

**(K)** $^{13}$C-pyruvate as in (J).

**(L)** $^{13}$C-R5P as in (J).

**(M)** $^{13}$C-E4P as in (J).

**(N)** $^{13}$C-citrate as in (J).

**(O)** $^{13}$C-G6P as in (E).

**(P)** $^{13}$C-F1,6-BP as in (E).

**(Q)** $^{13}$C-R5P as in (E).

**(R)** $^{13}$C-E4P as in (E).

**(S)** $^{13}$C-citrate as in (E).

**(T)** $^{13}$C-malate as in (E).

G6P, glucose-6-phosphate; F 1,6-BP, fructose 1,6-bisphosphate; R5P, ribose-5-phosphate; E4P, erythrose-4-phosphate; S7P, sedoheptulose-7-phosphate; S 1,7-BP, sedoheptulose 1,7-bisphosphate.

All data are represented as mean ± SEM from n=3 biological replicates.

*$p<0.05$, **$p<0.01$, ***$p<0.001$, ****$p<0.0001$ as determined by Two-Way ANOVA unless otherwise indicated.

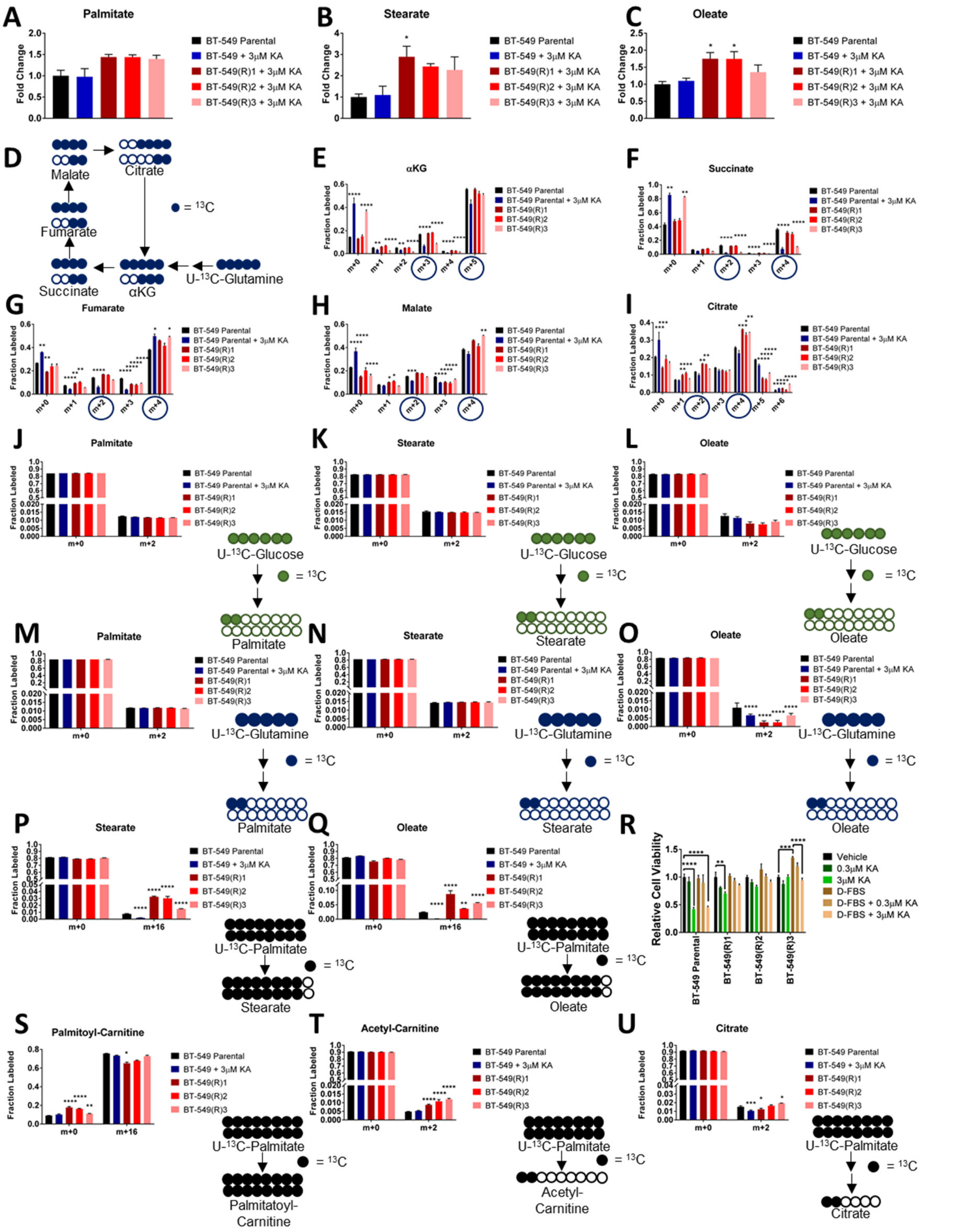


**Figure S4.** ***Related to Figure 4*****. Changes in fatty acid metabolism emerge as a functional output of evolved resistance to KA.**

**(A)** Levels of palmitate after treatment with vehicle or KA for 6 hours.
**(B)** Stearate as in (A).

**(C)** Oleate as in (A).
**(D)** Schematic depicting two turns of the TCA cycle after labeling cells with U-$^{13}$C-glutamine.
**(E)** Fraction labeled of $^{13}$C-αKG from U-$^{13}$C-glutamine labeling after pretreatment with vehicle or KA for 6 hours followed by tracing for 24 hours. Isotopomers labeled by U-$^{13}$C-glutamine are circled in blue.
**(F)** $^{13}$C-succinate as in (E).
**(G)** $^{13}$C-fumarate as in (E).
**(H)** $^{13}$C-malate as in (E).
**(I)** $^{13}$C-citrate as in (E).
**(J)** Fraction labeled of $^{13}$C-palmitate after pretreatment with vehicle or KA for 6 hours followed by U-$^{13}$C-glucose labeling for 24 hours.
**(K)** $^{13}$C-stearate as in (J).
**(L)** $^{13}$C-oleate as in (J).
**(M)** Fraction labeled of $^{13}$C-palmitate after pretreatment with vehicle or KA for 6 hours followed by U-$^{13}$C-glutamine labeling for 24 hours.
**(N)** $^{13}$C-stearate as in (M).
**(O)** $^{13}$C-oleate as in (M).
**(P)** Fractions labeled of $^{13}$C-stearate after pretreatment with vehicle or KA for 6 hours followed by U-$^{13}$C-palmitate labeling for 24 hours.
**(Q)** $^{13}$C-oleate as in (P).
**(R)** Cell viability of BT-549 parental and acquired resistant cells treated with vehicle or KA for 24 hours in complete media or media supplemented with 10% delipidated serum.
**(S)** Fraction labeled of $^{13}$C-palmitoyl-carnitine after pretreatment with vehicle or KA for 6 hours followed by U-$^{13}$C-palmitate labeling for 24 hours.
**(T)** $^{13}$C-acetyl-carnitine as in (S).
**(U)** $^{13}$C-citrate as in (S).

All data are represented as mean ± SEM from n=3 biological replicates.

*p<0.05, **p<0.01, ***p<0.001, ****p<0.0001 as determined by Two-Way ANOVA.

αKG, α-ketoglutarate.

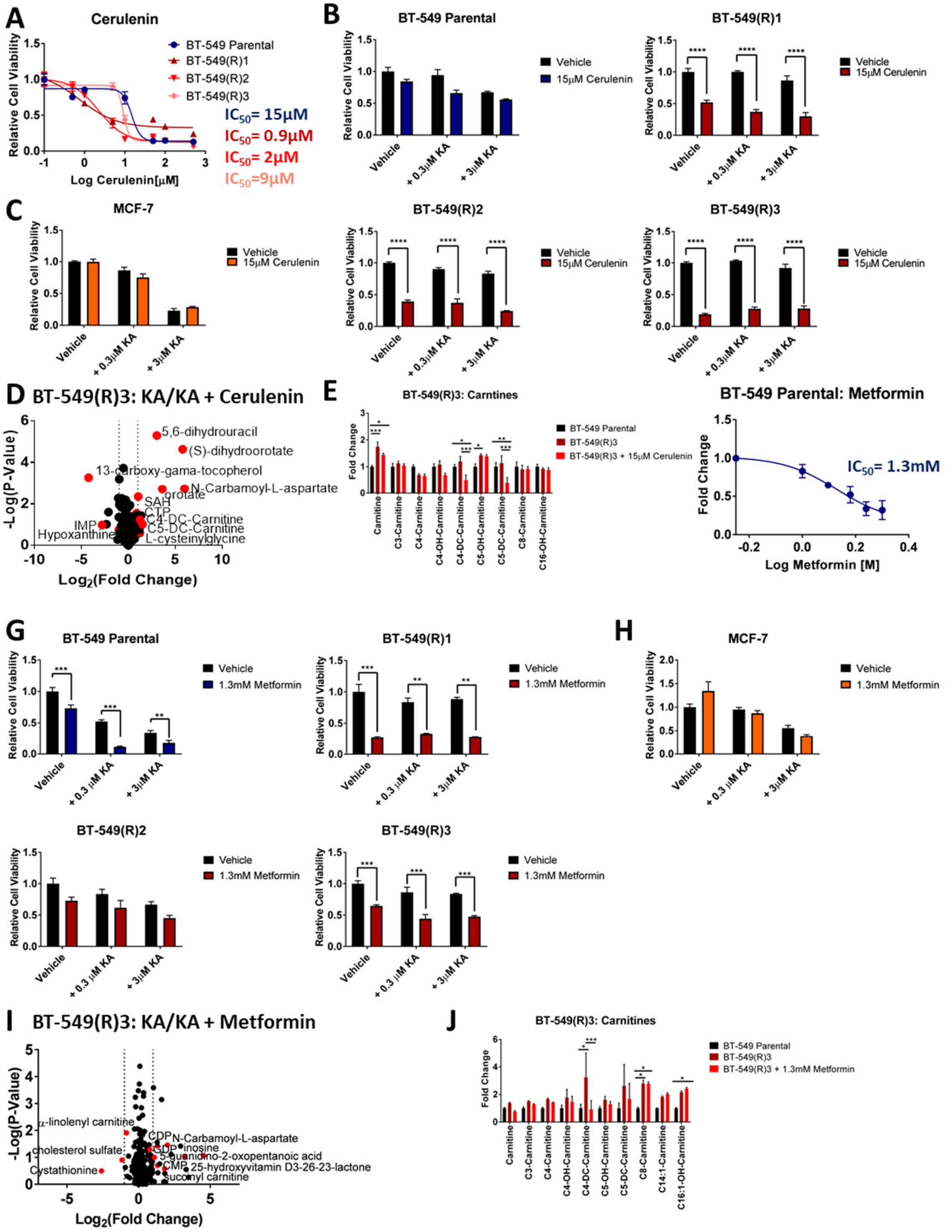


**Figure S5.** ***Related to Figure 5.*** **Changes in fatty acid metabolism occur downstream of differences in glycolysis in acquired resistance to KA.**

**(A)** BT-549 parental and acquired resistant cells dose-response curve treated with 0-500μM cerulenin for 24 hours.

**(B)** Cell viability of BT-549 parental cells and acquired resistant cells treated with cerulenin (15μM) with or without KA (0.3μM or 3μM) for 24 hours.

**(C)** Cell viability of MCF-7 cells as in (B).

**(D)** Volcano plot showing metabolite profiles of BT-549 acquired resistant (R)3 cells maintained in KA (3μM) with or without cerulenin (15μM). $Log_2$ fold change versus –$log_{10}$ p-value. Dotted lines along x-axis represent ± $log_2(1)$ fold change and dotted line along y-axis represents $-log_{10}(0.05)$. Metabolites ± $log_2(1)$ fold change shown as red points with metabolite names denoted. All other metabolites are black points.

**(E)** Acyl-carnitine levels in BT-549 acquired resistant (R)3 cells maintained in KA (3μM) and treated with or without cerulenin (15μM) for 6 hours.

**(F)** BT-549 parental cell dose-response curve treated with 0-2mM metformin for 24 hours.

**(G)** Cell viability of BT-549 parental and acquired resistant cells treated with metformin (1.3mM) with or without KA (0.3μM or 3μM) for 24 hours.

**(H)** Cell viability of MCF-7 cells as in (G).

**(I)** Volcano plot showing metabolite profiles of BT-549 acquired resistant (R)3 cells maintained in KA (3μM) with or without metformin (1.3mM) as in (D).

**(J)** Acyl-carnitine levels in BT-549 acquired resistant (R)3 cells maintained in KA (3μM) and treated with or without metformin (1.3mM) for 6 hours.

All data are represented as mean ± SEM from n=3 biological replicates.

*$p<0.05$, **$p<0.01$, ***$p<0.001$, ****$p<0.0001$ as determined by Two-Way ANOVA unless otherwise indicated.
